# Caltubin regulates microtubule stability via Ca^2+^-dependent mechanisms favouring neurite regrowth

**DOI:** 10.1101/2023.01.23.525163

**Authors:** Andrew Barszczyk, Julia Bandura, Qi Zhang, Haitao Wang, Marielle Deurloo, Yasmin Ahmed, Aiping Dong, Paul Meister, Jeffrey Lee, Hong-Shuo Sun, Yufeng Tong, Zhong-Ping Feng

## Abstract

Microtubule regulation is highly controlled in nerve regeneration. Caltubin, a novel Lymnaea stagnalis protein, contains putative EF-hand calcium-binding motifs and promotes neuronal outgrowth in Lymnaea and mouse. Here, we generated cell-permeable caltubin proteins to investigate mechanisms underlying this effect. We observed increased neurite extension and outgrowth following injury in caltubin-treated mouse neurons compared to vehicle controls. Purified caltubin bound α-tubulin between its L391-V405 amino acids and promoted microtubule assembly. Caltubin competitively inhibited binding of tubulin tyrosine ligase, which catalyzes tubulin retyrosination, and increased the ratio of detyrosinated to tyrosinated tubulin. Our crystal structure analysis confirmed that caltubin has four Ca^2+^-binding EF-hand motifs, like calmodulin but has distinct peptide binding domains. Our work suggests a unique Ca^2+^-dependent regulatory mechanism of microtubule assembly by caltubin. This may represent an essential mechanism of axonal regulation, which may optimize its activity in response to various calcium states, both physiological and following injury.

## Introduction

Central nervous system dysfunction, arising from a wide variety of disorders, including traumatic injury, neurodegenerative diseases, and many others, afflicts millions each year and constitutes a major source of disability and morbidity (Hyder, Wunderlich, Puvanachandra, Gururaj, & Kobusingye, 2007; Lilienfeld & Perl, 1994). It is well established that central neurons reside in inhibitory environments which are unfavorable for re-growth (Fawcett & Asher, 1999; Grandpré, Strittmatter, & Strittmatter, 2001; Mukhopadhyay, Doherty, Walsh, Crocker, & Filbin, 1994; Wang et al., 2002) and lack intrinsic growth capacity to do so (Sun & He, 2010). There are no existing therapies that enable the central neural regeneration after injury.

Cytoskeletal microtubules are a central component of cell structure (Ingber, 1993), intracellular transport (Gross, 2004), and intracellular signaling (Gundersen & Cook, 1999). Following injury, the tip of the neurite reseals and forms either a growth cone or a retraction bulb. A growth cone is characterized by bundled microtubules and is capable of regeneration, while a retraction bulb is characterized by disorganized microtubules and is incapable of regeneration (Bradke, Fawcett, & Spira, 2012). The mature central nervous system favors the formation of retraction bulbs because of a lack of intracellular growth cues and the presence of growth-inhibitory factors (Etienne-Manneville, 2010), as well as a lack of cytoskeletal dynamics that favour regrowth (Rodemer, Gallo, & Selzer, 2020). Microtubules are in a constant, dynamic equilibrium of assembly and disassembly with free α/β-tubulin heterodimers. These dynamics are tightly regulated by microtubule-binding proteins, which directly affect microtubule stability (Wade, 2009). Their recruitment to microtubules is controlled by a further set of enzymes, which post-translationally modify tubulins in response to the needs of the cell. Modifications like detyrosination and acetylation are associated with long-lived or “stable” microtubules (Moutin, Bosc, Peris, & Andrieux, 2021; Palazzo, Ackerman, & Gundersen, 2003; Webster, Gundersen, Bulinski, & Borisy, 1987) and changes in these modifications play an important role in promoting recovery from neuronal damage (Cho & Cavalli, 2012; Gobrecht et al., 2016). Second messengers like calcium are crucial for coordinating these molecular events (Schliwa, Euteneuer, Bulinski, & Izant, 1981), particularly in the growth cones of elongating neurons (Timothy M. Gomez & Zheng, 2006; Henley & Poo, 2004) and in the resealing of injured axon tips after injury (Rehder, Jensen, & Kater, 1992; Spira, Oren, Dormann, & Gitler, 2003). Microtubule stabilizer proteins, such as tau, MAP2, and MAP1B, promote axonal growth in healthy neurons (Chuckowree & Vickers, 2003; Harada et al., 1994; Sengottuvel, Leibinger, Pfreimer, Andreadaki, & Fischer, 2011; Sharma, Kress, & Shafit-Zagardo, 1994; Takei, Teng, Harada, & Hirokawa, 2000), direct intrinsically-driven neurite outgrowth (Bomze, Bulsara, Iskandar, Caroni, & Pate Skene, 2001; Leibinger et al., 2009; Leon, Yin, Nguyen, Irwin, & Benowitz, 2000), and desensitize neurite tips to growth-inhibitory factors (thus attenuating retraction bulb formation) (Erturk, Hellal, Enes, & Bradke, 2007). Microtubule-stabilizing agents promote regeneration of injured mouse nerves *in vivo* (Hellal et al., 2011; Sengottuvel et al., 2011). However, the mechanisms that underlie the regulation of microtubule stability remains unclear.

Our lab has identified a novel snail protein named caltubin (Nejatbakhsh et al., 2011), which is required for neuronal outgrowth and regeneration in snail neurons. When it is introduced into mouse neurons, caltubin also promotes neurite outgrowth and attenuates the retraction of lesioned neurites (Nejatbakhsh et al., 2011). The lack of a close caltubin homolog in the mouse genome, yet the conservation of its effect on neurite outgrowth of mouse neurons, suggests a conserved mechanism in both species. The sequence of caltubin contains several putative EF-hand like motifs, suggesting the possibility of calcium binding (T. M. Gomez, Robles, Poo, & Spitzer, 2001; Henley & Poo, 2004). In this work, we first produced a cell-penetrating version of the caltubin protein to enable its efficient delivery into mammalian cells. We then tested whether cell-penetrating caltubin functions to promote neurite outgrowth and neurite regrowth after injury in mammalian neuronal cell cultures, and determined 1) whether caltubin directly stabilizes microtubules, 2) whether caltubin competitively inhibits the function of any microtubule-regulating proteins, and 3) whether caltubin has the structural components necessary to allow for its regulation by calcium. In this way, we investigated whether the caltubin cell penetrating peptide serves as a microtubule stabilizer with the potential for endogenous regulation, which thus could constitute a novel therapeutic approach for repairing damaged central neurons.

## Results

### HIV-Tat peptide fusion delivers caltubin into live cells and promotes neurite outgrowth in primary neuronal culture

We produced a recombinant cell-penetrating version of caltubin to enable rapid, efficient and controlled delivery of caltubin into live cells. This cell-penetrating version of caltubin (CaT-Tat) consists of the full caltubin sequence fused with the protein transduction domain (PTD; residues 47-57 of HIV Tat protein (Frankel & Pabo, 1988; Vives, Brodin, & Lebleu, 1997)) on the C-terminus end and a thrombin-cleavable His6 tag affixed to the N-terminus (see Supplemental Figure 1A for full sequence). The His6 tag facilitates protein purification from *E. coli* and the PTD peptide enables CaT-Tat to traverse cell membranes. Protein expression and purity was confirmed via western blot (Supplemental Figure 1B). Cell penetration of CaT-Tat was confirmed in mouse hippocampal culture, where His6 tag immunoreactivity is apparent in both cell bodies and neurites (Supplemental Figure 1C). To determine whether CaT-Tat retains the neurite outgrowth-promoting function of transfected caltubin in mammalian cells (Nejatbakhsh et al., 2011), primary mouse cortical culture at a day-in-vitro (DIV) 2 was treated with CaT-Tat or vehicle control for 24 hours. Cells were then fixed, permeabilized and immunostained for β-tubulin to allow for visualization and quantification of neurite morphology using fluorescence microscopy. Mouse cortical cultures treated with caltubin exhibited statistically significant increases in total neurite outgrowth (331.3 ± 18.3 μm; n=90) compared to those treated with vehicle (247.3 ± 12.4 μm; n=143). The longest neurites, which frequently but not certainly are fated to become axons as development continues, were also significantly longer in cells treated with CaT-Tat (198.7 ± 13.8 μm; n=90) compared to cells treated with vehicle alone (108.9 ± 7.0 μm; n=143) (Figure 1). There was no apparent effect on the number of branches between CaT-Tat-treated (7.0 ± 0.4; n=90) and vehicle-treated (6.8 ± 0.3; n=143) cells.

**Figure 1.**
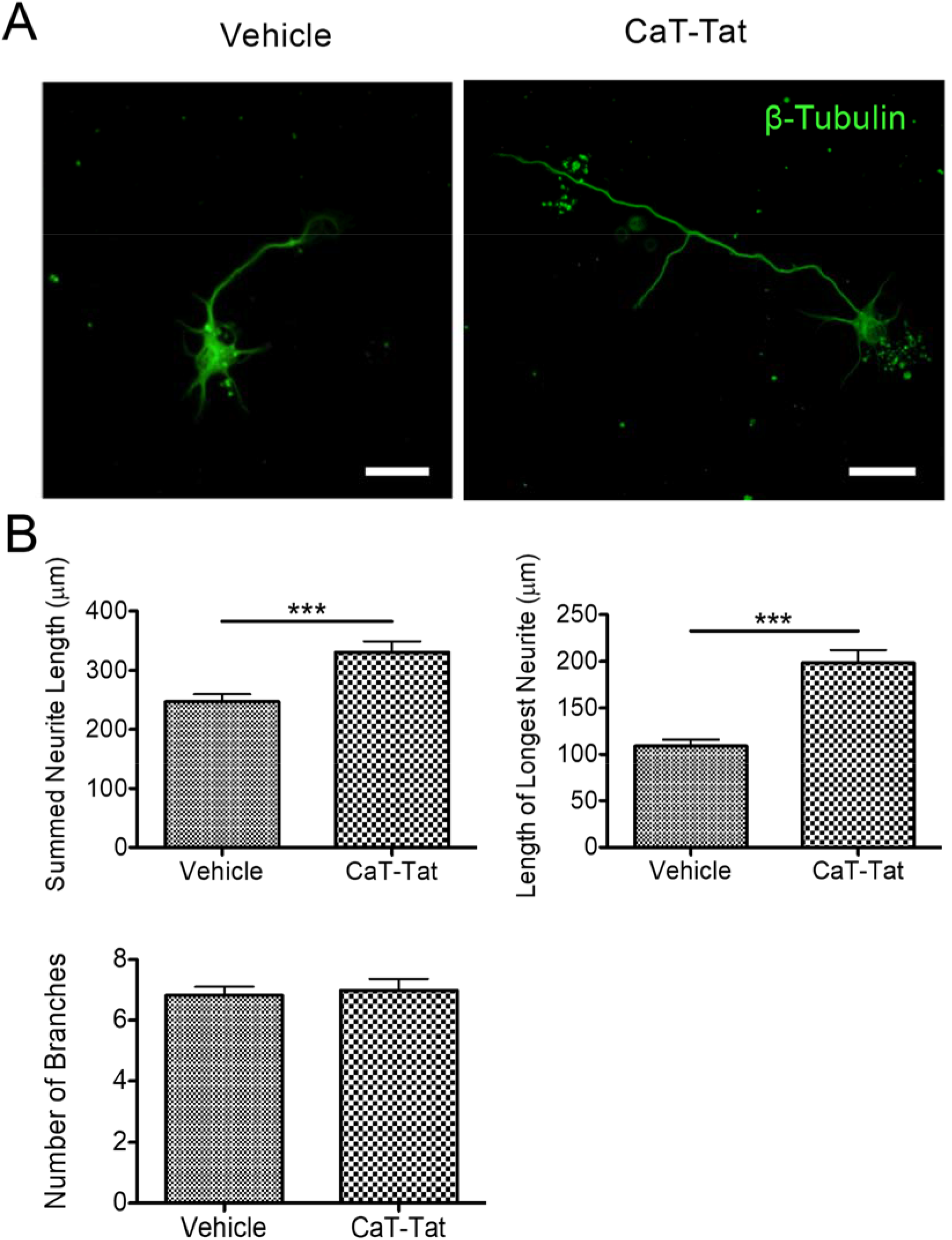
Caltubin-Tat (CaT-Tat) increases neurite outgrowth in primary mouse cortical neurons. (A) Representative images of primary mouse cortical culture treated with CaT-Tat (1 μM) or vehicle (PBS) on DIV 2. Cells were fixed and permeabilized on DIV 3, and then immunolabeled with β-tubulin antibody and visualized with fluorescence microscopy. Scale bar = 20 μm. (B) Characterization of mouse cortical cell morphology after treatment with PBS vehicle (n=143) or CaT-Tat (n=90) from 3 independent experiments. Mean ± SEM; *** p < 0.05 in 2-way Student’s t-test.

### Caltubin enables re-growth of damaged neurites in primary neuronal culture

To determine whether CaT-Tat regulates neurite outgrowth or retraction in injured neurites, we pre-treated mouse cortical cultures with 1 μM CaT-Tat or FITC-LC-Tat control peptide for one hour, lesioned them, and then quantified neurite length at 2-minute intervals for 14 minutes. In neurons pretreated with control peptide, lesioned neurites retracted an average of 18.8 ± 1.8 μm (n=5) from the site of the lesion over the 14-minute time window (Figure 2). Conversely, neurites in CaT-Tat-treated cells elongated an average of 24.4 ± 3.2 μm (n=4).

**Figure 2.**
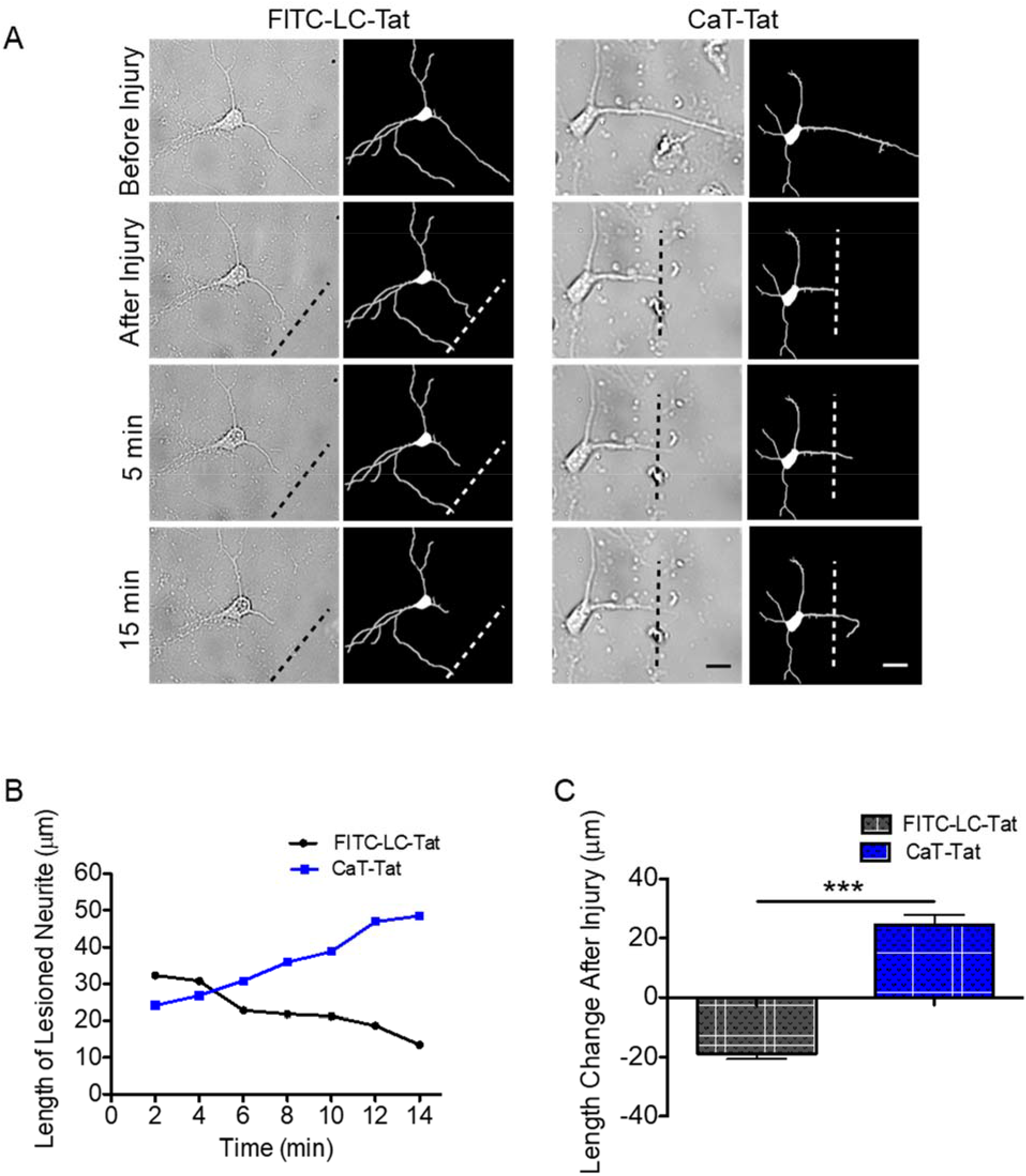
Caltubin-Tat (CaT-Tat) enables neurite outgrowth in axotomized neurites. Mouse cortical neurons were pretreated with 1 μM FITC-LC-Tat or 1 μM CaT-Tat for one hour, mechanically lesioned at approximately two cell body diameters from the cell body and then observed for 15 minutes. (A) Images from single neurons taken at regular intervals, with the lesion site marked by a dotted line. A tracing of neurites accompanies each image to aid visualization of the extent of each neurite. Scale bar represents 21 μm. (B) Quantification of neurite length of the lesioned neurite depicted in panel A. (C) Mean length change of lesioned neurites after 15 minutes in cells treated with FITC-LC-Tat (n=5) or CaT-Tat (n=4). Mean ± SEM; *** p < 0.0001 in Student’s t-test.

### Caltubin interacts with the C-terminal region of α-tubulin

The pro-growth functionality of endogenous caltubin in snail (Nejatbakhsh et al., 2011) and the present study’s demonstration of the same effect of CaT-Tat on mouse neurons implies that caltubin may manipulate a conserved mechanism. We previously proposed that caltubin function is mediated by tubulins (Nejatbakhsh & Feng, 2011; Nejatbakhsh et al., 2011), which are highly conserved between snails and mammals, including rodents. Here, we investigated the details of this interaction to gain a better understanding of how caltubin may function via tubulins. We first determined whether CaT-Tat binds directly to tubulins by performing an affinity pull-down assay with α/β-tubulin complex purified from bovine brains. Purified tubulin protein was used in place of cell homogenate to reduce the potential for false-positive binding to tubulin via intermediary adaptor proteins. We found that α-tubulins within this tubulin mixture interacted with GST-caltubin, with no appreciable binding to GST alone (Supplemental Figure 2).

To determine where caltubin binds to tubulin, peptide arrays comprising overlapping 15 amino acid segments of α-tubulin (TUBA1) and β-tubulin (TUBB3) were constructed, and then probed with caltubin to determine where it binds to either isoform. TUBA1 represents a common isoform of α-tubulin, and TUBB3 is a neuron-specific isoform of β-tubulin. Upon probing these arrays with His6-tagged CaT-Tat, several tubulin segments on each array exhibited His6 tag immunoreactivity (Figure 3A; indicated by teal or red arrows). Blue arrows indicate His6 peptide and were used as positive controls. At this point, three of the most likely caltubin binding regions were selected based on signal intensity and their exposure in the 3D structure of tubulins. These were L391 (TUBA1 array), C211 (TUBB3 array) and E431 (TUBB3 array) - all three are indicated with red arrows on the array (Figure 3A).

**Figure 3.**
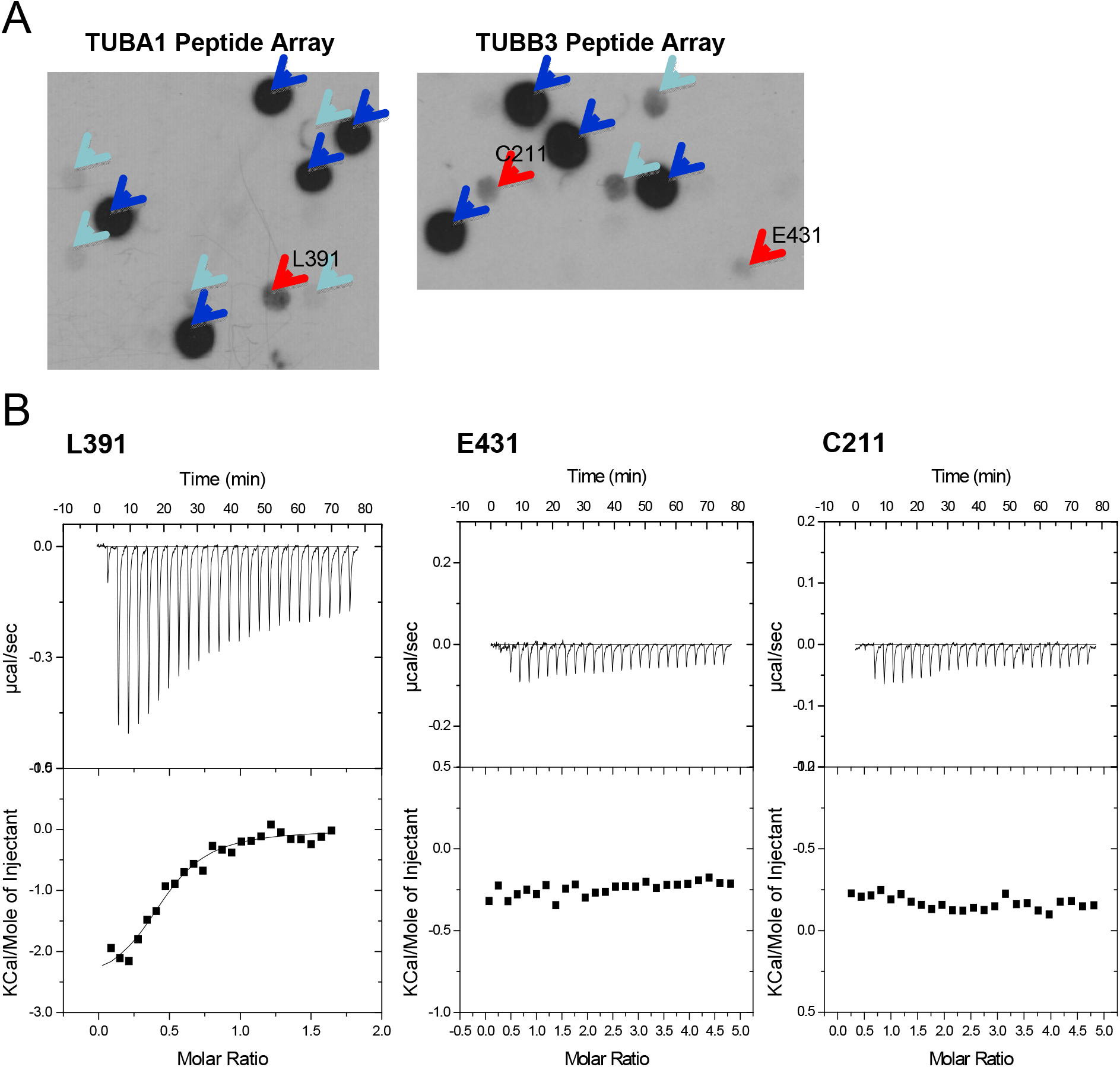
Identification of a His-caltubin-Tat (CaT-Tat) binding domain on TUBA1. (A) TUBA1 and TUBB3 peptide arrays were probed with His-CaT-Tat (1 μM), with detection of the His6 signal. Dark blue arrows indicate positive controls (HHHHHHHAA), light blue arrows indicate segments where significant signal was detected, and red arrows indicate binding regions that were subsequently tested with ITC. (B) Isothermal titration calorimetry of L391 (LDHKFDLMYAKRAFV; Kd = 3.42 μM), E431 (EEEGEMYEDDEEESE) or C211 (CFRTLKLATPTYGDL) peptides into a 1μM CaT-Tat solution.

To confirm the interaction of the peptides with caltubin, we used isothermal titration calorimetry (ITC) to quantify binding affinity. Only one of the three candidate peptides – the ‘L391’ peptide within the C-terminal region of TUBA1 (peptide sequence: LDHKFDLMYAKRAFV) – exhibited an appreciable energy change upon titration of CaT-Tat (*K_D_* = 3.42 μM) (Figure 3B), demonstrating binding to caltubin. Together, these data demonstrate that caltubin binds α-tubulin at its L391 region directly.

### Caltubin promotes assembly of microtubules from purified tubulin in vitro

The ability of caltubin to directly bind purified tubulin suggests that it may directly affect microtubule assembly. Therefore, we probed the effect of caltubin on the rate and extent of spontaneous assembly of tubulin heterodimers into microtubules using an *in vitro* microtubule assembly assay. Our data demonstrate that in this assay, microtubules assembled spontaneously in the presence of either CaT-Tat or vehicle control (Figure 4A). However, the maximum reaction velocity, in OD/minute, was 60% greater in the presence of CaT-Tat than in the presence of vehicle control: the slopes of the reaction curves during this phase were .0097 ± .0002 (n=3) and .0061 ± .0001 (n=3), respectively (Figure 4B). Further, there were 37.5% more assembled microtubules in the CaT-Tat condition than in the vehicle condition upon reaction equilibrium (.1432 ± .0007 versus .1969 ± .0012, respectively; Figure 4C). Therefore, CaT-Tat promotes both the rate and extent of tubulin assembly into microtubules, even in a system devoid of microtubule-associated proteins.

**Figure 4.**
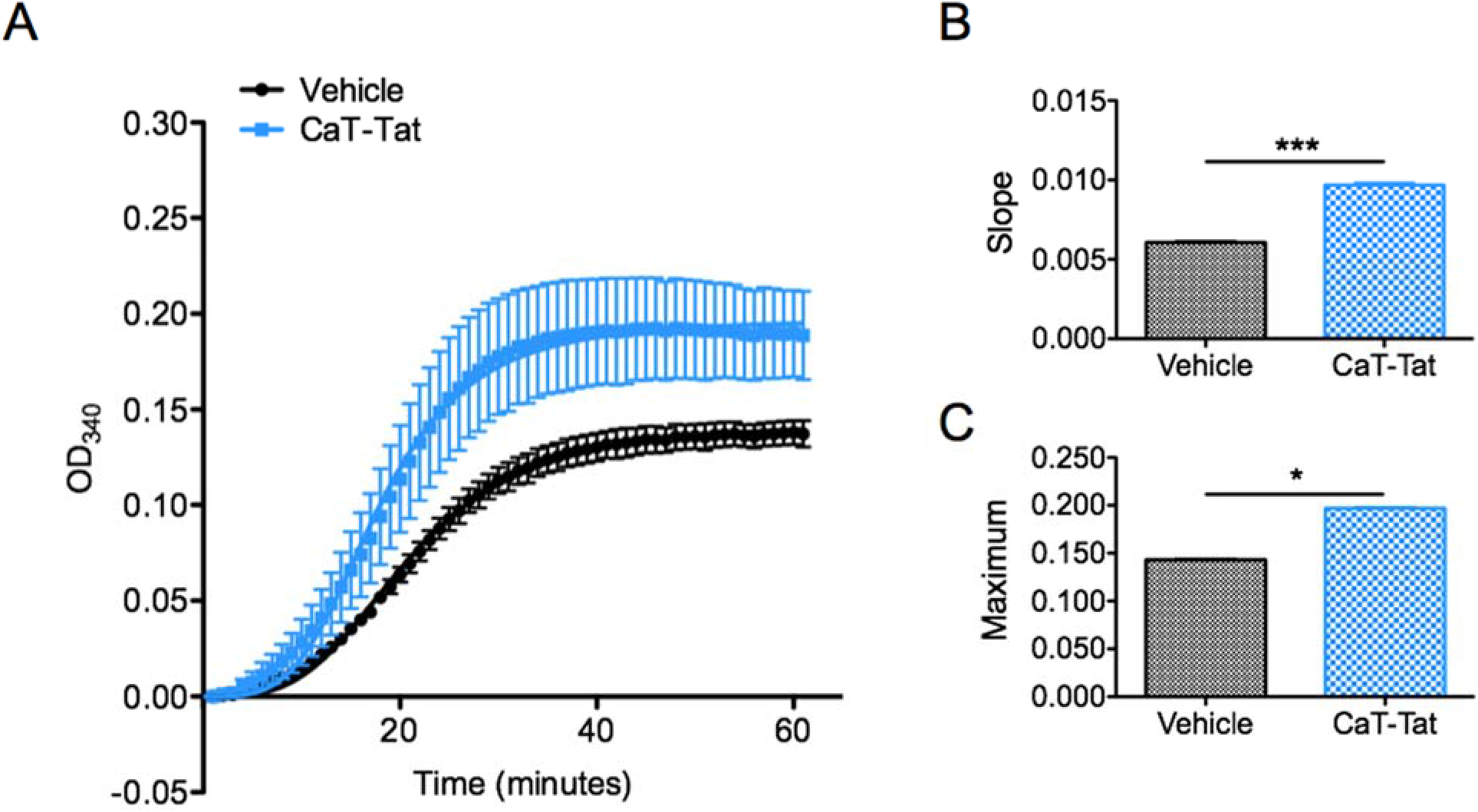
Caltubin-Tat (CaT-Tat) enhances the assembly of purified porcine tubulin into microtubules in vitro. (A) Tubulin polymerization indicated by optical density (OD, at 340 nm; normalized to T=0) change over time in tubulin-containing wells treated with CaT-Tat (24 μM) or vehicle buffer (80 mM PIPES, pH 7.3, 2 mM MgCl2, 50 mM NaCl, 1 mM DTT, 100 mM L-arginine, 1 mM EGTA). Points represent mean ± SEM from 3 experiments, with data fit to an allosteric sigmoidal function (solid lines). (B) Linear regression was used to calculate the slope (maximum reaction velocity, in OD/minute) between 16 and 23 minutes (r^2^ = .9984 for vehicle, r^2^= .9986 for CaT-Tat). The slope for the vehicle line is .006063 +/.000098 (n=3) and the slope for the CaT-Tat line is .009679 +/- .000147 (n=3). Therefore, CaT-Tat causes a 60% increase in general assembly velocity, which is statistically significant by t-test (p<0.0001). (C) To quantify the extent of the reaction, the curve was fit to an allosteric sigmoidal function (indicated by sigmoidal curve; r^2^ = .9984 for vehicle, r^2^= .9960 for CaT-Tat). The calculated peak value was .1432 +/- .0007 for the vehicle and .1969 +/- .001243 for CaT-Tat, equating to a 37.5% increase in the extent of microtubule polymerization. There is no overlap between the 95% confidence intervals of .1416-.1477 for vehicle and .1944-.1994 for CaT-Tat.

### Caltubin displaces tubulin tyrosine ligase from tubulins

Though caltubin was able to promote microtubule assembly independently of microtubule-regulatory proteins, we also tested the hypothesis that caltubin functions by interacting with such proteins. Mapping of the L391 peptide to the 3D structure of tubulin in complex with tubulin tyrosine ligase (TTL; PDB: 402L) revealed that binding to the L391 peptide region would likely interfere with the interaction of interaction of TTL with tubulin heterodimers (Figure 5A). We thus proceeded to investigate whether caltubin might interfere with TTL function, using a co-immunoprecipitation assay. Specifically, TTL from mouse cortical tissue was immunoprecipitated and the amount of TTL-associated tubulin that immunoprecipitated along with it was probed. In the presence of CaT-Tat, less tubulin precipitated out of solution with TTL (Figure 5B), even when normalized to the quantity of TTL (Figure 5C). When an excess of L391 peptide was added to the cell lysate to occupy the caltubin binding site for tubulin, CaT-Tat no longer reduced the immunoprecipitation of tubulin by TTL (Figure 5D). These data show that caltubin competitively inhibits binding of TTL to tubulin by binding to α-tubulin in the L391-V405 region.

**Figure 5.**
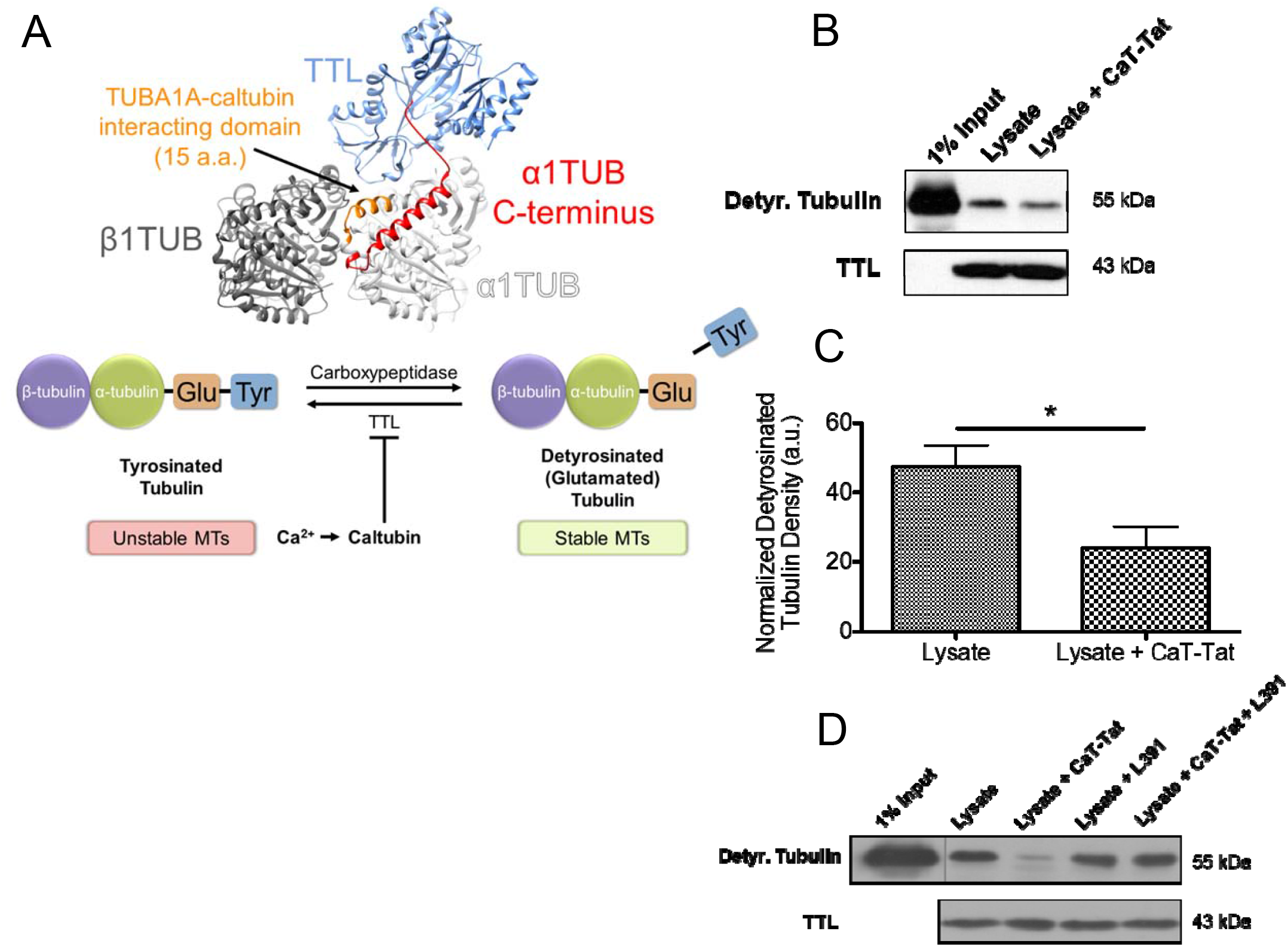
Tubulin displaces TTL from microtubules. A) The apparent close proximity of His-caltubin-Tat to the TTL binding site may prevent the interaction of TTL with free α-tubulins (top right panel; PDB ID: 4IHJ (Prota et al., 2013), created with UCSF Chimera (Pettersen et al., 2004)) and thus inhibit their retyrosination, resulting in a greater ratio of detyrosinated to tyrosinated tubulin (bottom panel). B) Co-immunoprecipitation of detyrosinated tubulin with tubulin tyrosine ligase (TTL) from mouse cortical tissue lysate in the presence of no additives, .42 μM CaT-Tat, 0.42 μM L391 peptide (LDHKFDLMYAKRAFV) or both His-caltubin-Tat and L391 peptides. Lysis/Wash Buffer: 50 mM Tris, 150 mM NaCl, 1% NP-40, 0.25% Sodium deoxycholate, pH 7.4. After all immunoprecipitations, proteins were eluted, separated using SDS-PAGE and probed for detyrosinated tubulin. (B) Western blot running 20 μg of protein per lane. Detyrosinated tubulin was detected under basal conditions or in the presence of CaT-Tat (C) Densitometry quantifying the immunoprecipitation of detyrosinated/tyrosinated tubulin from the above blot. Detyrosinated tubulin signal density was normalized to the 1% input condition within each experiment. n = 4 independent experiments. * p < 0.05 by Student’s t-test. (D) Single experiment conducted as above, but with the addition of the L391 peptide as a possible inhibitor of caltubin function.

### Caltubin affects tyrosination of tubulin, a marker of microtubule stability

TTL is an enzyme which regulates the tyrosination state of microtubules by adding a tyrosine at the C-terminus of detyrosinated α-tubulin (Ersfeld et al., 1993; Janke & Kneussel, 2010; Prota et al., 2013). Interference with TTL function by caltubin should therefore translate to a decrease in microtubule tyrosination. We used an immunocytochemical approach to test the effects of caltubin treatment on tubulin tyrosination state. Furthermore, to confirm that this inhibition translates into functional effects, we probed the effect of caltubin on neurite outgrowth and recovery from injury.

In the first experiment, cell-penetrating caltubin (Cat-Tat) was applied onto DIV 1 primary mouse cortical culture for 24 hours to determine whether it affects microtubule stability in cells exhibiting neuronal morphology. Microtubule stability was assessed immunocytochemically by quantifying the ratio of immunoreactivity for detyrosinated tubulin (Figure 6A), which is associated with stable/long-lived microtubules, to that of tyrosinated tubulin, which is associated with unstable microtubules (Gundersen, Kalnoski, & Bulinski, 1984). Mouse cortical cells treated with caltubin exhibited a greater ratio of detyrosinated/tyrosinated tubulin relative to vehicle-treated cells. When considering the detyrosinated/tyrosinated ratio as a function of distance from the soma, this difference was apparent throughout most of the cell, including the soma (Figure 6B; n = 54-72 from 3 experiments). When the mean overall ratio of detyrosinated/tyrosinated tubulin was compared between vehicle-treated and CaT-Tat treated cells (Figure 6C; n = 54-72 from 3 experiments), CaT-Tat treated cells exhibited a higher detyrosinated/tyrosinated tubulin ratio (2.3 ± 0.14; n=72) than vehicle-treated cells (1.11 ± 0.08; n = 54). A significant difference also persisted when the ratio in only the distal-most 5 μm was considered (Figure 6D), with CaT- Tat treated cells exhibited a higher ratio of detyrosinated to tyrosinated tubulin (1.497 ± 0.107; n=72) than vehicle-treated cells (0.998 ± 0.119; n = 54). Finally, a separate experiment was done to exclude the possibility that the Tat PTD was mediating these effects. In this experiment, CaT-Tat treated cells exhibited a higher detyrosinated/tyrosinated tubulin ratio (0.980 ± 0.037; n=75) than FITC-LC-Tat (0.682 ± 0.036; n = 48) or vehicle-treated cells (0.679 ± 0.025; n = 66). There was no statistically significant difference between the ratios of vehicle-treated cells and FITC-LC-Tat-treated cells (Figure 6E). These data therefore suggest that caltubin stabilizes neuronal microtubules and may thus regulate neurite outgrowth and regrowth.

**Figure 6.**
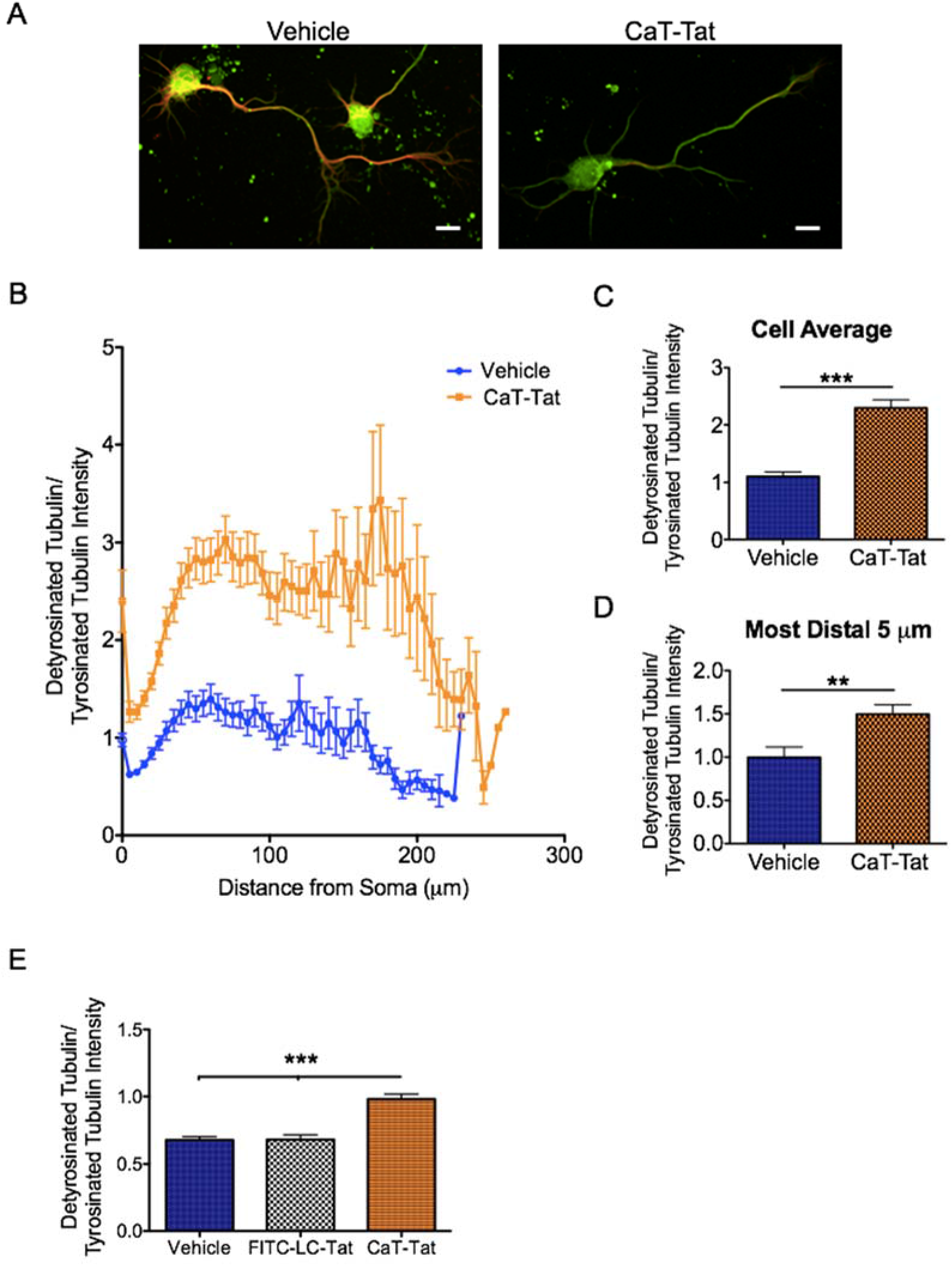
CaT-Tat increases the ratio of detyrosinated (stable) to tyrosinated (unstable) tubulin in cultured cortical cells. (A) Representative images of primary mouse cortical cells treated with vehicle control (phosphate-buffered saline; PBS) or 1 μM CaT-Tat for 24 hours. Green: detyrosinated (stable) tubulin marker; Red: tyrosinated (unstable) tubulin marker. Scale bar = 10 μm. (B) Ratio of detyrosinated to tyrosinated tubulin in neurites as a function of distance from the soma of mouse cortical cells treated with vehicle (PBS) or 1 μM CaT-Tat for 24 hours. Cells with longest neurite length greater than 2 SD above the mean were considered outliers and excluded (2 cells per group). (C) Mean ratio of detyrosinated to tyrosinated tubulin within mouse cortical cells treated with vehicle control or 1 μM CaT-Tat for 24 hours. (D) Mean ratio of detyrosinated to tyrosinated tubulin in the most distal 5 μm segment of mouse cortical cells treated with vehicle control or 1 μM CaT-Tat for 24 hours. Mean ± SEM. ** p < 0.01 and *** p < 0.001 in two-tailed t-test. n = 54-72 from 3 experiments. (E) Mean ratio of detyrosinated to tyrosinated tubulin within mouse cortical cells treated with vehicle control, 1 μM FITC- LC-Tat or 1 μM CaT-Tat for 48 hours. *** p < .0001 in one-way ANOVA with Bonferroni correction. n = 48-75 cells.

### Crystal structure of caltubin reveals a calmodulin-like fold

We have previously shown by sequence analysis that caltubin contains several EF-hand motifs that are known to bind calcium (Nejatbakhsh et al., 2011). However, the exact number of EF-hands and the overall structure of caltubin remain elusive. We solved the structure of caltubin using X-ray crystallography. Caltubin was over-expressed in *E. coli* as a polyhistidine-SUMO-fusion protein and the tag-removed protein was purified to homogeneity and set up for crystallization trials. Caltubin only crystallized in conditions that contain calcium salt; however, the crystals did not diffract to a high resolution. We optimized the diffraction quality of the crystals in the presence of strontium chloride (SrCl2) and solved the crystal structure of the core domain of caltubin (a.a. 36–179) using single-wavelength anomalous diffraction signal from strontium ions in space group *P2*_1_ at 1.3 Å resolution. Each asymmetric unit contains two copies of the molecule with a pseudo-*C2* non-crystallographic symmetry.

Each molecule is bowl-shaped and the N-terminus of one molecule occupies the central cavity of the other molecule (Figure 7A). The two copies of caltubin in the asymmetric unit have an r.m.s.d. of 0.24 Å over all atoms, suggesting near-identical conformations. The caltubin core domain comprises eight α-helices that can be grouped into four helix-loop-helix motifs, each form an EF-hand in complex with one Sr^2+^ ion. The four Sr^2+^ ions are exposed and located in the outer surface of the bowl shape. The first two and the last two EF-hands are clustered into two groups. In each group, the two EF-hands are packed in a head-to-head orientation with a short β-type interaction between the two loops from the EF-hands. Interestingly, the second EF-hand (hereinafter referred to as EF2) contains a 13 a.a. motif that coordinates a Sr^2+^ ion, whereas the other three contains the canonical 12 a.a. motif as seen in most other EF-hands (Figure 7B). Of the three canonical EF-hands (EF1, EF3, EF4), 5 or 6 of the 7 coordinating atoms are provided by the protein (Figure 7C) and the rest are provided by either water molecules or ethane-1,2-diol from the crystallization buffer, whereas for EF2, all 7 atoms coordinating the Sr^2+^ ion are provided by the protein. An unusual proline P99 before the last coordinating glutamic acid (E100) adopts a less common *cis conformation* that facilitates the coordination of Sr^2+^ (Figure 7D). All other prolines in the structure are in a *trans* conformation. Caltubin is a very acidic protein with a calculated isoelectronic point (pI) of 4.8 for the core domain, and 5.0 for the full-length. This is consistent with the electrostatic potential map (Figure 7E) which shows dominantly negative electrostatic potentials on the surface. The crystal structure also shows several other Sr^2+^ ions, which do not seem to be play a structural role.

**Figure 7.**
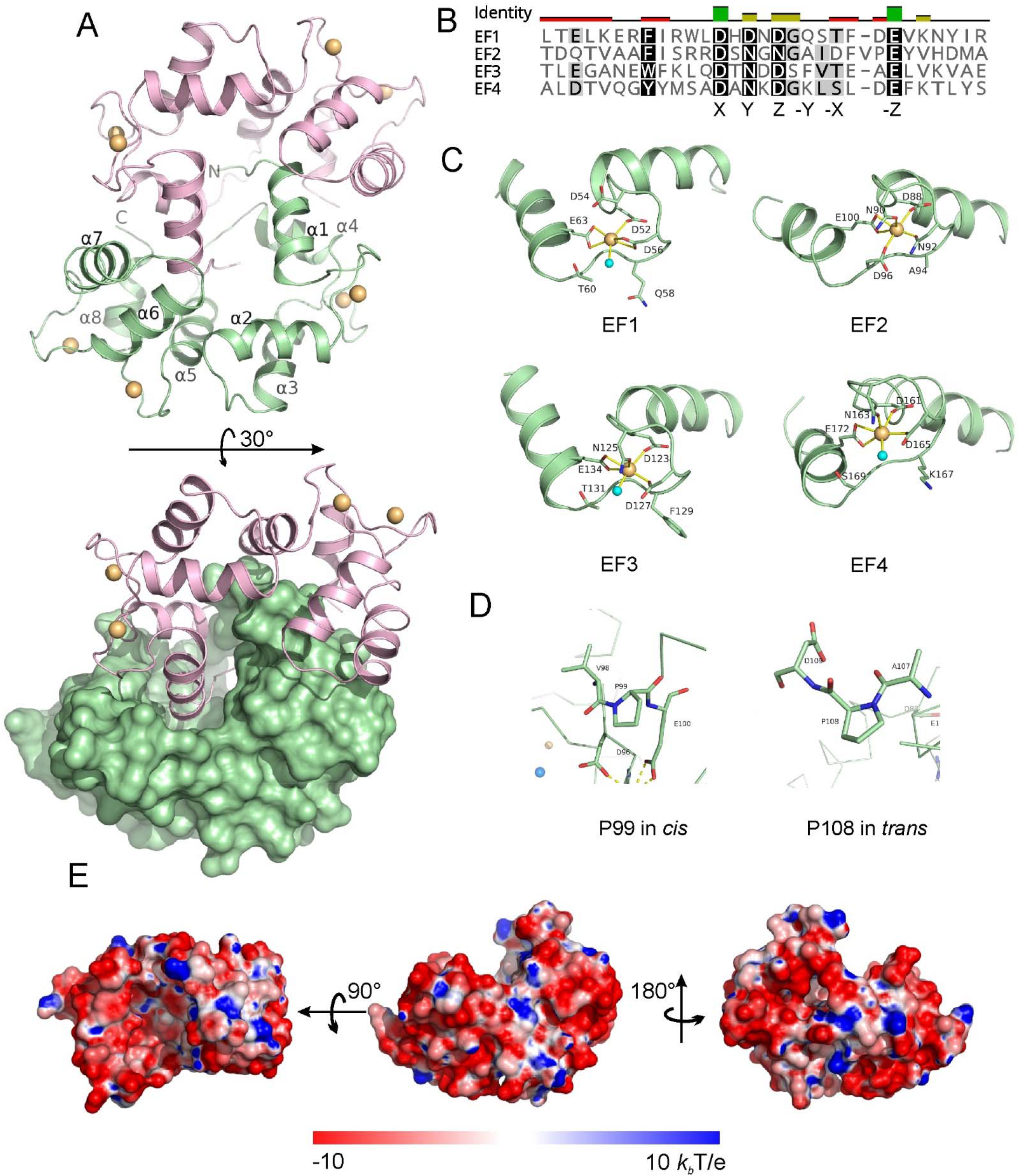
Crystal structure of caltubin. (A) Each asymmetry unit contains two caltubin molecules. Each caltubin comprise 8 α-helices that form 4 EF-hands. Structural Sr^2+^ ions are shown in orange. Non-structural Sr^2+^ ions are omitted. (B) Sequence alignment of the 4 EF-hands of caltubin. EF2 is one residue longer than the other 3 EF-hand calcium binding loops. X, Y, Z, -X, -Y, -Z indicates residues that are normally involved in coordinating the bound metal ions. (C) Details of the coordination of the Sr2+ in the four EF-hands. (D) The proline P99 in EF2 adopts an uncommon cis conformation, whereas other prolines in the molecule adopt a trans conformation, P108 is shown as an example for comparison. (E) Surface electrostatic potential of caltubin calculated by PDB 2PQR server (Baker, Sept, Joseph, Holst, & McCammon, 2001) using the default parameters and shown at ± 10 Kbit/e scale, where kb is the Boltzmann’s constant, T is the temperature in K, e is the charge of an electron.

A Dali search revealed the closest structure homolog of caltubin is the calcium-bound calmodulin in complex with the IQ motif peptide of voltage-gated calcium channel Ca_V_1 (PDB: 2VAY) with a Z-score of 14.6 and an overall r.m.s.d of 3.2 Å for all the residues. However, structural overlay of the first two EF hands of caltubin with those of calmodulin shows an r.m.s.d. of 1.5 Å, and the last two EF-hands an r.m.s.d. of 0.9 Å (Figure 8A), significantly smaller than the overall r.m.s.d. This suggests the caltubin can also be divided into two subdomains, like calmodulin. It is well established that the hinge region between the N- and C-terminal subdomains of calmodulin is critical for the conformational flexibility required for binding target proteins and for the regulation by calcium ions. Interestingly, the equivalent region in caltubin is 3 a.a. shorter than that of calmodulin (Figure 8B) and contains a proline P108, a known α-helix breaker. This suggests that despite that similarity of the structural fold of caltubin with that of calmodulin, the regulation of molecular recognition events of the two proteins with effector proteins will be vastly different, which is to be tested in future studies.

**Figure 8.**
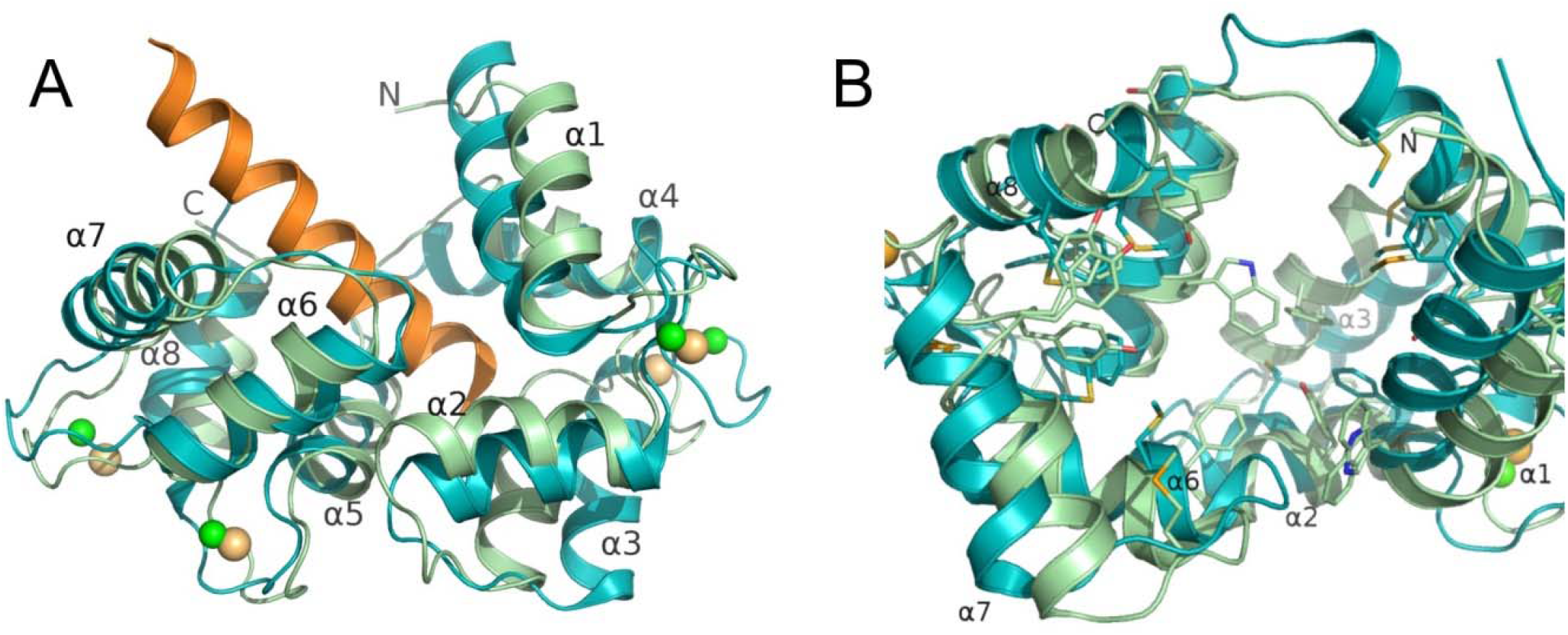
Comparison of caltubin and calmodulin structures. (A) Overlay of the structure of caltubin (green) with that of calmodulin (cyan) in complex with CaV IQ motif (orange, PDB 2VAY). Sr2+ ions are shown in light-orange spheres, and Ca2+ ions in green. (B) Aromatic residues (Tyr, Phe, Trp) and methionine in the peptide binding pocket. Caltubin has only two methionines in the putative peptide binding pocket whereas it is enriched with aromatic residues. Some aromatic residues of caltubin have alternative conformations.

## Discussion

Axonal regeneration is a key solution to central nerve injury. We aimed to identify critical molecules responsible for enhancing capacity for nerve repair. In this study, we delivered a cell-penetrating caltubin protein into mammalian neurons and found that it can promote neurite extension and the regeneration of calmodulin. The second EF-hand of caltubin (EF2) has a novel 13-a.a motif and an unusual proline in *cis* injured neurites. By solving the crystal structure of caltubin, we confirmed caltubin as an EF-hand type calcium-binding protein with a structure similar to calmodulin. We further showed that caltubin binds to α-tubulin directly to promote assembly of microtubules. Caltubin interaction with α-tubulin interferes with tubulin tyrosine ligase and alters the tyrosination/detyrosination balance of microtubules in neurons. This is the first study to demonstrate the fast-acting effect of directly delivered caltubin protein on promoting axonal regeneration via microtubule regulation compared to ectopically expressed caltubin.

We determined that caltubin promotes microtubule assembly in an assay containing highly pure tubulin (with a near-absence of MAPs). This suggests that caltubin works directly on microtubules to stabilize them (e.g. against depolymerizing processes or agents (Takemura et al., 1992)) and/or to promote favorable conformations of tubulin heterodimers or protofilaments to facilitate their integration into the microtubule. These processes shift the equilibrium of microtubule assembly in the positive direction. Thus, in situations within adult central neurons when microtubule stability is suboptimal for neurite outgrowth (Arimura & Kaibuchi, 2007), caltubin could promote microtubule assembly and thus promote outgrowth. Microtubule stabilizers are well known to promote outgrowth in this way (Esmaeli-Azad, McCarty, & Feinstein, 1994; Sengottuvel et al., 2011), so long as they remain at an optimal concentration (Derry, Wilson, & Jordan, 1995; Derry, Wilson, Khan, Ludueña, & Jordan, 1997). Our data suggest that caltubin may act similarly to microtubule stabilizers to promote neurite outgrowth by directly stabilizing microtubules.

To determine microtubule regulator-dependent mechanisms of caltubin function, we identified a caltubin-binding site on the C-terminal α-helix of α-tubulin (L391-V405) and then investigated whether caltubin interferes with microtubule-regulatory mechanisms in this vicinity. We found that caltubin blocks the association of the tubulin tyrosine ligase (TTL) enzyme with tubulin. TTL preferentially associates with non-polymerized α-tubulin in the free tubulin pool (Szyk, Deaconescu, Piszczek, & Roll-Mecak, 2011; Wehland & Weber, 1987) and re-tyrosinates it (Argaraña, Arce, Barra, & Caputto, 1977; Kumar & Flavin, 1981). Indeed, our findings show that caltubin causes an increase in the proportion of detyrosinated versus tyrosinated tubulin in microtubules. Therefore, these results show that caltubin may promote neurite outgrowth by interfering with tubulin tyrosine ligase (TTL) activity and affecting microtubule tyrosination/detyrosination balance. It remains unclear how caltubin-induced increases in detyrosinated tubulin that occur in the interchangeable tubulin pool would become incorporated into the microtubule. The conventional view is that tubulin joins the microtubule in its tyrosinated state and becomes detyrosinated over time (Webster et al., 1987). If caltubin indeed functions by regulating TTL activity on free tubulins, our findings would imply that detyrosinated tubulin can be incorporated into microtubules as well. Further study is needed to reconcile this point.

Recent findings implicate TTL in the regulation of neurite outgrowth and regeneration, adding plausibility to the notion that caltubin regulates these cell behaviors by interfering with TTL function. TTL-null cortical and hippocampal neurons have been shown to exhibit longer neurites, resist microtubule depolymerization by nocodazole, and block CLIP-170 association with neurite tips and growth cones (Erck et al., 2005). Similarly, in cerebral granule neurons, increasing microtubule tyrosination reduces neurite outgrowth (Inami, Omura, Kubota, & Konishi, 2018), suggesting that inhibiting TTL, and therefore microtubule tyrosination, may be permissive for neurite outgrowth. Our data, which show that caltubin causes an increase in cortical neurite outgrowth over a 1-2 day period (Figure 1), are consistent with the morphological consequences of a TTL knockout as well as decreases in microtubule tyrosination and support the notion that caltubin functions by interfering with TTL activity. If this is so, our study further supports the involvement of TTL and microtubule tyrosination in regulating neuronal morphology.

Interestingly, disruption of TTL function and decreases in tyrosination does not lead to neurite extension in all neurons. In one study, TTL knockout in DRG neurons markedly reduced neurite outgrowth (Barnat et al., 2016), which was consistent with an inhibitory effect of increased detyrosination on neurite extension in these neurons (Gobrecht et al., 2016). Additionally, the chemogenetic activation of DRG neurons, leading to the enhancement of axonal regrowth, has been shown to be associated with increased tyrosinated tubulin (Wu et al., 2020), suggesting that increased tyrosination may favour regeneration in certain contexts. Thus, the role of TTL across cell types and developmental stages should be studied in more detail.

Our data demonstrate that caltubin, in addition to its neurite extension-promoting effects in normal neurons, promotes outgrowth following neurite lesion in an *in vitro* cortical neuronal model. Given these findings, the microtubule polymerization-promoting effects of caltubin take on additional significance in neurite injury. Following lesioning, the injured neurite tip must restructure into a growth cone rather than a retraction bulb in order for regeneration to occur, and this process is contingent on microtubule dynamics (Bradke et al., 2012; Cajal & May, 1991). In fact, following injury *in vivo*, administration of nocodazole promotes formation of retraction bulbs while paclitaxel favours formation of growth cones (Erturk et al., 2007). A significant portion of this early phase of regeneration occurs within minutes of a neurite lesion (Nejatbakhsh et al., 2011). While the structure of injured neurite terminals was not analyzed in this study, the occurrence of early regeneration means that caltubin-dependent remodeling of the neurite tip into a growth cone has necessarily taken place. Therefore, similarly to paclitaxel, caltubin may stabilize microtubules and hinder the formation of a retraction bulb, leading to its demonstrated positive effect on recovery from injury.

Based on sequence alignment, we previously proposed that caltubin is a putative EF-hand calcium binding protein (Nejatbakhsh et al., 2011). However, the number of EF-hand motifs was not confirmed. In this study, our crystal structure unequivocally confirmed that caltubin is a calmodulin-like protein, containing 4 EF-hands which bind Sr^2+^. Sr^2+^ has similar cation size, coordination geometry, and hydration energies as those of Ca^2+^, and was used in this study to obtain the high-resolution structure of caltubin. The Sr^2+^ ion in all four EF-hands showed pentagonal bipyramidal geometry (Fig. 7). It is worth noting that the second EF-hand (EF2) of caltubin has a 13 a.a. calcium-binding loop with a single-residue insertion in the C-terminal part of the loop. Of all known EF-hand domains of known crystal structure, the only other EF-hand containing a 13 a.a. calcium-binding loop is the EF1 of extracellular protein osteronectin (Hohenester et al., 1996; Hohenester, Sasaki, Giudici, Farndale, & Bächinger, 2008). All other known non-canonical EF-hands with normal Ca^2+^ coordination contain insertions or deletions are located in the N-terminal part of the loop (Gifford, Walsh, & Vogel, 2007; Grabarek, 2006). Furthermore, the calcium-binding loop of caltubin EF2 contains an unusual Pro99 in *cis* conformation that facilitates the coordination of the Sr^2+^ by: the main chain carbonyl group of Ala94, the side chain carbonyl group of Asn90 and Asn92, the bident carboxyl side chain of Glu100, and one carboxyl group from each of the side chain of Asp96 and Asp98. Unlike the Sr^2+^ ion in the other three EF-hands of caltubin, where small molecule ligands (water or ethane-1,2-diol) contributed to the coordination, the seven atoms that coordinate with Sr^2+^ in EF2 were all provided by the protein. Typically, the contribution of solvent entropy plays an important role in the preference of EF-hands of binding Ca^2+^ over Mg^2+^, another divalent cation present at a high concentration in the cell. Therefore, whether caltubin has a higher affinity towards Mg^2+^ at EF2 due to lack of solvent-mediated coordination remains to be investigated. The 13-a.a. EF-hand motif in osteronectin is important for the sequence-specific collagen recognition (Hohenester et al., 2008). Similarly, the unusual sequence and conformation of EF2 may also be important for the pro-growth function of caltubin.

Caltubin being a calcium-binding protein, and therefore subject to calcium regulation, has several implications for its usefulness as a promoter of microtubule stability and subsequently neurite outgrowth and regeneration. The overstabilization of microtubules by microtubule stabilizers like paclitaxel poses a problem for the usefulness of all known microtubule-stabilizing drugs – namely that unlike endogenous proteins, their function is not subject to appropriate regulation either in expression or protein activity (Baas & Ahmad, 2013). Caltubin may be unique in this respect because it has evolved in snails to manipulate evolutionarily conserved microtubule regulatory mechanisms and is likely to be subject to regulation by calcium, which plays major roles in regulating endogenous neurite outgrowth and cellular responses following injury. Therefore, it is possible that caltubin function can be controlled, like intracellular calcium signals, both spatially (i.e. preferentially activated in subcellular compartments where it is needed) and in response to cellular events (i.e. massive calcium fluxes in response to injury). In injury, as discussed previously, caltubin could therefore regulate the binding of proteins to microtubules in a calcium-dependent manner, resulting in “as-needed” changes in microtubule stability and protein transport. Since caltubin likely functions downstream of many mechanisms of microtubule regulation, yet also remains subject to cellular regulation (unlike small molecule regulators), it constitutes a highly dynamic regulator of microtubules. Nevertheless, the degree to which caltubin action is regulated by intracellular calcium signaling is not yet known and is an avenue for future study.

## Methods

All procedures were performed in accordance with the Canadian Council on Animal Care guidelines and approved by the Faculty of Medicine and Pharmacy Animal Care Committee at the University of Toronto.

### Cloning, Protein Overexpression, and Purification of Caltubin

The cDNA encoding the full-length caltubin was codon optimized for *E. coli* expression and synthesized (BioBasic Inc.). The construct encoding a cell penetrating caltubin that contained an N-terminal thrombin-cleavable His6 (hexahistidine) tag and a C-terminal 11 a.a. HIV-Tat protein transduction motif (YGRKKRRQRRR) was subcloned into a pET-46 Ek/LIC vector (EMD Millipore #71335). The construct encoding a GST-tagged caltubin was subcloned into a pGEX-4T-1 vector (Amersham #28954549). All plasmids were confirmed with restriction enzyme digestion and sequencing.

Plasmids were transformed into *E. coli* strain BL21(DE3) grown in lysogeny broth, in the presence of appropriate antibiotics, and overexpression of the proteins was induced by 0.1–0.5 mM isopropyl β-D-1-thiogalactopyranoside (IPTG, Sigma Aldrich, #I5502) for 20 hrs at 18°C for His6-tagged caltubin or 3 hrs at 30°C for GST-tagged caltubin, before being harvested by centrifugation.

For His6-tagged caltubin, harvested cell pellets were resuspended in lysis buffer (50 mM Tris-HCl, pH 8, 300 mM NaCl, 20 mM imidazole) supplemented with protease inhibitor cocktail (cat. PIC002.1, BioShop) and lysed using a benchtop cell disruption unit (Constant Systems Ltd.). Affinity chromatography (HIS-Select^™^ Nickel Affinity Gel; #P-6611; Sigma Aldrich Inc.) was then used to purify His6-tagged caltubin from the cell lysate and the protein was eluted with 125 mM imidazole (in lysis buffer). Finally, pooled eluent was dialyzed 1:1000 against phosphate-buffered saline (PBS; #21600- 069; Life Technologies Inc.) 3 times at 4°C in regenerated cellulose dialysis tubing (MWCO=3.5 kDa). SDS-PAGE was followed by anti-His6 Western blot to confirm protein expression and Coomassie Brilliant Blue staining to assess the purity of the protein. Finally, the protein was concentrated to the desired concentration using centrifugal filters (#UFC800308; Amicon^®^ Ultra centrifugal filters, Ultracel^®^ −3K or larger; Merck Millipore Ltd.)

GST fusion protein was similarly purified using affinity chromatography by incubating bacterial cell lysate with glutathione-Sepharose 4B beads (#17075601; GE Healthcare) according to the manufacturer’s instructions and eluted from beads with elution buffer (50 mM Tris-HCl, pH 8, 20mM reduced glutathione, 5% glycerol).

For crystal structure determination, the DNA encoding the caltubin core domain (residues 37-179) was subcloned in a pET28-MKH8SUMO vector (for TEV protease cleavable N-terminal His8-SUMO tag). The protein was over-expressed in a phage-resistant BL21 (DE3) *E. coli* strain harbouring a pRARE2 plasmid (Agilent) using Terrific Broth cultured in a LEX Bioreactor (Epiphyte3). Protein production was induced using 0.5 mM IPTG for overnight growth at 17 °C. The proteins were first purified using nickel-nitrilotriacetic acid (NTA) agarose resin (Qiagen), and the His8-SUMO tag was then removed by TEV protease (10:1 w/w) during overnight dialysis against a buffer containing 50 mM TrisHCl, pH 8.0, 150 mM NaCl, 3 mM β-mecaptoethanol, and 1 mM CaCl2. Uncleaved proteins, SUMO tag, and TEV protease were removed by another pass through the nickel-NTA resin. The proteins were further purified using anion-exchange chromatography (Source 15Q, GE Healthcare) using a salt gradient from 0 to 1 M NaCl in a buffer containing 20 mM TrisHCl, pH 8.0, 1 mM DTT. The purified proteins were concentrated to a final concentration of 32 mg/mL, and stored at −80 °C. All protein concentrations were determined by measuring UV absorbance at 280 nm using NanoDrop 2000 (Thermo Fisher Scientific) and the molecular weight determined by mass spectrometer. The selenomethionine-derivative of caltubin was overexpressed in *E. coli* using a prepacked M9 SeMet growth media kit following the manufacturer’s instruction (Medicilon).

### Dissociated Neuronal Culture, Neuronal Injury, and Immunocytochemical Procedures

All procedures were performed in accordance with the Canadian Council on Animal Care guidelines and approved by the Faculty of Medicine and Pharmacy Animal Care Committee at the University of Toronto. Cortical and hippocampal neurons were isolated from embryonic day 17 (E17) pups of timed pregnant female CD1 mice (Charles River Laboratories (Quebec, Canada)). Mice were sacrificed by cervical dislocation and fetal cortices and hippocampi were harvested, digested with .025% trypsin with EDTA (Gibco) for 15 minutes at 37°C, then triturated with a pipette to homogeneity. Cells were filtered through a 70 μm cell strainer, centrifuged, and resuspended in plating Neurobasal medium (#21103049; Invitrogen) containing 1x penicillin/streptomycin (#P4333; Sigma Aldrich), 1x GlutaMAX^™^ (#35050061; Thermo Fisher Scientific), 10% fetal bovine serum (#F1051; Sigma Aldrich). 4.5 x 10^4^ dissociated cells in 60 μl of plating medium were plated on 12 mm coverslips (#194310012; Bellco) pre-coated with 0.1 mg/ml poly-D-lysine hydrobromide solution (#P0899; Sigma), and cultured at 37°C and 5% CO2. After 3 hours, the plating solution was replaced with 450 μl of culture medium (Neurobasal, 1x penicillin/streptomycin; 1x GlutaMAX^™^; 2% B27 supplement (#17504044; Thermo Fisher Scientific).

In order to assess effects of caltubin treatment on recovery from injury, primary mouse cortical cells were lesioned on DIV 2 on their longest neurite (presumed axon) at two cell body lengths (~20 μm) from the cell body. Phase contrast microscopy was then used to image the neuron every two minutes for 16 minutes, and changes in the length of the proximal segment of the lesioned neurite were quantified.

To characterize the effects of caltubin treatment on tyrosination/detyrosination balance of microtubules, immunocytochemistry was performed as previously described (Barszczyk et al., 2015). Briefly, cells on coverslips were fixed with 4% paraformaldehyde/1x PBS, washed, and then permeabilized/blocked with 0.1% Triton^™^ X-100 (#T8787; Sigma)/1% bovine serum albumin (BSA; #ALB001.5; BioShop Canada Inc.)/1x PBS. Cells were then immunolabeled with primary antibodies (see below) for 2 hours at room temperature, washed, and immunolabeled with the corresponding secondary antibody for 30 minutes at room temperature. Coverslips were washed and mounted on glass slides using ProLong^®^ Gold Antifade Mountant (#P36930; Thermo Fisher Scientific). Antibodies used for immunocytochemistry were: mouse anti-β-tubulin (1:1000 immunocytochemistry; 1:500 immunohistochemistry) (#T0198, Sigma), goat anti-mouse-488 (1:500) (#A11001; Invitrogen); rabbit anti-detyrosinated (glutamated) tubulin (1:200) (#AB3201, Millipore), goat anti-rabbit-405 (1:500) (#A31556; Invitrogen); rat anti-tyrosinated tubulin (1:1000) (#AB6160, Abcam), goat anti-rat-568 (1:500) (#A11077; Invitrogen); Mouse anti-His6 (1:100; 1h for immunocytochemistry) (#sc-8036; Santa Cruz Biotechnology); goat anti-mouse-488 (1:200) (#A11001; Invitrogen). Immunolabeled cells were imaged on a Zeiss LSM 700 laser confocal microscope (Carl Zeiss Inc.). Pinhole size was adjusted for cell or tissue thickness and reduced when imaging a single plane. The experimenter was blinded to treatment condition prior to imaging, and un-blinded upon completion of data analysis.

### Analysis of Neuronal Morphology and Posttranslational Modifications Obtained from Immunocytochemistry

Neurons were selected in phase contrast images of live cells based on the following criteria: (1) rounded cell body, (2) presence of neurites longer than the diameter of the cell body, (3) absence of blebbing, and (4) no overlap with other neurites significant enough to obscure the boundaries of all neurites. Neurites were traced semi-automatically using the NeuronJ plug-in (ver. 1.4.2; Erik Meijering) for ImageJ (ver. 1.46a; National Institutes of Health, USA) to determine total neurite length, the length of the longest neurite and the number of neuritic branches per cell.

Immunofluorescence of detyrosinated versus tyrosinated tubulin in immunolabeled cells was analyzed using the SynD plugin (Schmitz et al., 2011) for MATLAB^®^ (Mathworks). Intensity readings for each tubulin type as a function of distance from the soma was analyzed to obtain a ratio of detyrosinated to tyrosinated tubulin immunoreactivity at any given distance from the soma. For average readings across an entire cell, the intensities at each distance from the cell were averaged to get an average intensity reading for the entire cell.

### Peptide Array Generation and Probe

The peptide array was synthesized using the SPOT technique(Frank, 2002) using a MultiPep RS peptide synthesizer (Intavis AG) on a modified cellulose membrane following manufacturer’s instructions. For quantification of the binding of tubulin peptides with caltubin, three α-tubulin peptides, L391 (LDHKFDLMYAKRAFV), E431(EEEGEMYEDDEEESE), and C211 (CFRTLKLATPTYGDL) were chemically synthesized by Genscript (USA) (HPLC purified, >90% purity). Each peptide is N-acetylated and C-amidated to reduce the effect of charged terminal residues. Fluorescein isothiocyanate (FITC)-labeled Tat peptide (FITC-LC-Tat, a.a. 47-57) was purchased from AnaSpec, Inc. (Toronto, Canada). The binding assay was performed as previously described (Nady *et al*, 2008). The array was blocked with 5% skim milk in PBS/T (1x PBS + 0.1% Tween20) overnight at 4°C, washed, and then probed with 1 μM His-caltubin-Tat in PBS overnight at 4°C with shaking. After washing, His-caltubin-Tat was detected with mouse anti-His6/goat anti-mouse IgG-HRP antibodies and then visualized with an ECL kit (3 min reaction time; #E2500; Denville Scientific Inc.). Antibodies used for array were: mouse anti-His6 (1:1000) (#sc-8036; Santa Cruz Biotechnology; 4°C overnight), anti-mouse IgG-HRP (1:4000; 1h at room temperature, #AP130P; Chemicon International).

### GST-Pulldown and TTL Immunoprecipitation

Cortical tissue lysate was obtained from adult female CD1 mice. Briefly, cortical tissue was washed and then lysed in lysis buffer (50 mM Tris, pH 7.4, 150 mM NaCl, 1% Nonidet^®^ P40 (#NON505; BioShop Canada Inc.), 0.25% sodium deoxycholate) including protease inhibitor cocktail (1 mM PMSF (#PMS123.5), 1 μg/mL aprotinin (#APR600.10), 1 μg/mL leupeptin (LEU001.5), 1 μg/mL pepstatin A (#PEP605.5)), all from BioShop Canada Inc.), mechanically triturated, and vortexed to promote solubilization. Samples were centrifuged and supernatants used for subsequent experiments.

To confirm caltubin binding to α-tubulin using affinity pull-downs involving GST-tagged caltubin, 20 μl of glutathione-Sepharose 4B beads were pre-incubated with 500 μg of tissue lysate (9.2 μg/ul) in lysis buffer (PBS, 0.2% Triton X-100, 1 mM PMSF, 1 μg/mL aprotinin, 1 μg/mL leupeptin, 1 μg/mL pepstatin A) for 1h at 4°C and then briefly washed. Beads were then incubated with a mixture of 2 μg/μl of tissue and 50 μg of GST fusion protein in PBS for 2h at 4°C. Beads were then washed, and proteins were eluted from the beads (elution buffer: 10 mM reduced glutathione/50 mM Tris-HCL pH 8.0/5% glycerol). Eluted proteins were separated using SDS-PAGE, transferred to a membrane, and then probed with antibodies against α-tubulin, as described below.

To determine if caltubin interferes with the co-immunoprecipitation of specific tubulins with TTL, His-caltubin-Tat was added at 0.42 μM to 500 μl of cortical lysate (2 μg of tissue/μl of lysate), and 0.42 μM tubulin peptide L391 was added to displace caltubin from tubulin. This reaction mixture was incubated for 1h at 4°C, before adding 2 μg of anti-tubulin tyrosine ligase (TTL) antibody (2 μg/500 μL; ID3 immunoglobin (IgG); generously provided by Dr. Christian Erck (Erck et al., 2005; Wehland & Weber, 1987)) and incubating overnight at 4°C with agitation. The reaction mixtures were then incubated with 20 μl of pre-washed protein A/G beads (Santa Cruz Biotechnology sc-2003 for 1h at 4°C to immunoprecipitate the anti-TTL antibody and attached protein complexes. The beads were then washed, centrifuged, supernatant was removed, and protein complexes were eluted from the beads (elution buffer: 80 μl lysis buffer +20 μl SDS loading buffer). Proteins were then separated on a 10% SDS-PAGE, transferred to a membrane and then probed with antibodies against detyrosinated and tyrosinated tubulin, as described below.

### Western Blot

Protein concentration of samples was determined by the Pierce^™^ BCA Protein Assay Kit (#23227; Thermo Scientific). Equal amounts of sample protein (6-20 μg) were separated in an 8-15% SDS-PAGE gel and transferred to a nitrocellulose membrane (#66485; PALL Life Sciences) via semi-dry transfer (200 mA per gel, 60 min). Membranes were blocked with 5% skim milk (#SKI400.500, BioShop^®^ Canada Inc.) in Tris-buffered saline with 0.1% Tween-20 (TBST) and immunoblotted with primary antibodies in 1% milk/TBS overnight at 4°C with agitation. After washing with TBST, membranes were incubated with the corresponding HRP- or AP-tagged secondary antibodies for 1.5 h at room temperature with agitation. An ECL+ reagent (3 min; #E2500; Denville Scientific Inc.) was then used to visualize horseradish peroxidise (HRP)-tagged antibodies using chemiluminescence according to the manufacturer’s instructions. Alternatively, alkaline phosphatase conjugated antibodies were detected colourometrically using BCIP/NBT (nitro-blue tetrazolium and 5- bromo-4-chloro-3’-indolyphosphate) Substrate Solution (#BE116.100ML; Bio Basic Canada Inc). For analysis of bands, densitometry was employed to quantify proteins of interest using the gel analyzer function of ImageJ (ver. 1.46a; National Institutes of Health, USA). Antibodies used for western blot were: mouse anti-His6 (1:1000) (#sc-8036; Santa Cruz Biotechnology), goat anti-mouse IgG-AP (1:1000 in 5% BSA; #sc-2008; Santa Cruz Biotechnology, Inc.); mouse anti- alpha-tubulin (1:6000, #T6074; Sigma), anti-mouse IgG-HRP (1:6000; 2h at room temperature, #AP130P; Chemicon International); rabbit anti-detyrosinated (glutamated) tubulin (1:1500 overnight at 4°C) (#AB3201, Millipore), anti-rabbit-IgG-HRP (1:7500; 2h at room temperature, #7074S; Cell Signaling Technologies); rat anti-tyrosinated tubulin (1:1500 overnight at 4°C) (#AB6160, Abcam), donkey anti-rat-IgG-HRP (1:7500 2h at room temperature, #712035153; Jackson ImmunoResearch Inc.); mouse anti-TTL (1:2000); anti-mouse IgG-HRP (1:5000; 2h at room temperature, #AP130P; Chemicon International).

### Isothermal Titration Calorimetry (ITC)

His-caltubin-Tat was dialyzed into low-salt buffer 25 mM Tris-HCl, pH 8.0, 150 mM NaCl, 1 mM TCEP and concentrated with a centrifugal concentrator (Millipore) to 50 μM. The interaction of His- caltubin-Tat with 3 tubulin peptides (L391, E431, and C211) or CaCl_2_ was measured – each was solubilized in low-salt buffer to a final concentration of 1 mM for addition into the reaction vessel. ITC measurements were performed at 25°C on a VP-ITC Micro Calorimeter (Malvern). A total of 25 injections, each of 10 μL peptide or calcium solution, were delivered into a sample cell of 1.4 mL containing His-caltubin-Tat. The data was analyzed using the Origin for ITC software on the instrument and fitted to a one-site binding model.

### *In Vitro* Microtubule Assembly Assay

The high throughput screen (HTS) tubulin polymerization assay kit (Cat. BKOO4P, Cytoskeleton Inc.) was carried out as per the manufacturer’s instructions, with some minor modifications. Briefly, a reaction vessel was prepared on ice containing 3 mg/ml of >97% purified porcine tubulin (Cytoskeleton, Inc. (#T240)) and the substances being tested: vehicle or 24 μM His-caltubin-Tat. Both substances had been predialyzed into reaction buffer: 80 mM PIPES (PB0719-100G; Bio Basic), pH 7.3, 2 mM MgCl2 (#MAG510.500, BioShop), 50 mM NaCl, 1 mM DTT (#D9779; Sigma), 100 mM L-arginine (#A5131; Sigma), 1 mM EGTA (#E4378, Sigma). To start the reaction, 1 mM GTP (#R0461; Thermo Fisher Scientific) was added and the reaction volume was transferred to a pre-heated (37 dC) ½ area 96-well microplate (#3697, Corning) and OD340 readings were taken every minute for 60 minutes to obtain a time course of microtubule assembly.

### Crystallization, data collection and structure determination

Caltubin crystal was grown at 20°C using the sitting drop vaporization method by mixing 0.5 μL protein solution (32 mg/mL) with 0.5 μL well solution containing 24% PEG3350, pH 7.5 and 150 mM SrCl2. Harvested crystal was cryoprotected by immersion in the reservoir solution supplemented with 10% ethylene glycol before flash-frozen in liquid nitrogen. X-ray diffraction data were collected at the Canadian Light Source beamline 08ID-1. All datasets were processed with the HKL-2000 suite (Minor, Cymborowski, Otwinowski, & Chruszcz, 2006) and indexed in the *P*21 space group. The phase was solved using the single-wavelength anomalous diffraction (SAD) signal of strontium using the SOLVE/RESOLVE program (Terwilliger, 2003). The graphics program COOT(Emsley & Cowtan, 2004) was used for model building and visualization, REFMAC (v.5.8) (Murshudov, Vagin, & Dodson, 1997) for restrained refinement. The final models were validated by MolProbity(Chen et al., 2010). The crystal structure of caltubin is deposited in the protein databank as PDB ID code: 6VAN. All structural analyses and the preparation of molecular graphics were performed using PyMOL (Schrödinger, LLC) and UCSF Chimera (Pettersen et al., 2004).

### Statistics

Two-group comparisons were made by two-tailed Student’s t-test and multiple comparisons were made by one-way analysis of variance (ANOVA; Bonferroni correction for multiple comparisons) using GraphPad software (Prism).*p* < 0.05 were considered statistically significant.

## Acknowledgments

Support for this research was provided by the following funding agencies: Natural Sciences and Engineering Research Council of Canada (NSERC- RGPIN-2016-04574) to HSS, Canadian Institutes of Health Research (CIHR- PJT- 153155) to ZPF and YT. AB & JB were recipients of Ontario Graduate Scholarship (Doctoral). We would also like to acknowledge Jonathan Cook and Farshad Azimi for their technical advice regarding protein purification during this project, and Lin Pei for his technical support.

## Author contributions

AB and ZPF conceptualized the study. AB, QZ, HW, MD, YA, AD, PM, JL performed experiments and analyzed the data. AB, JB, HSS, YT, and ZPF prepared the manuscript.

## Materials & Correspondence

Correspondence and materials requests should be addressed to Dr. Zhong-Ping Feng.

## Supplementary Information

### Supplementary Figures

**Supplementary Figure 1.**
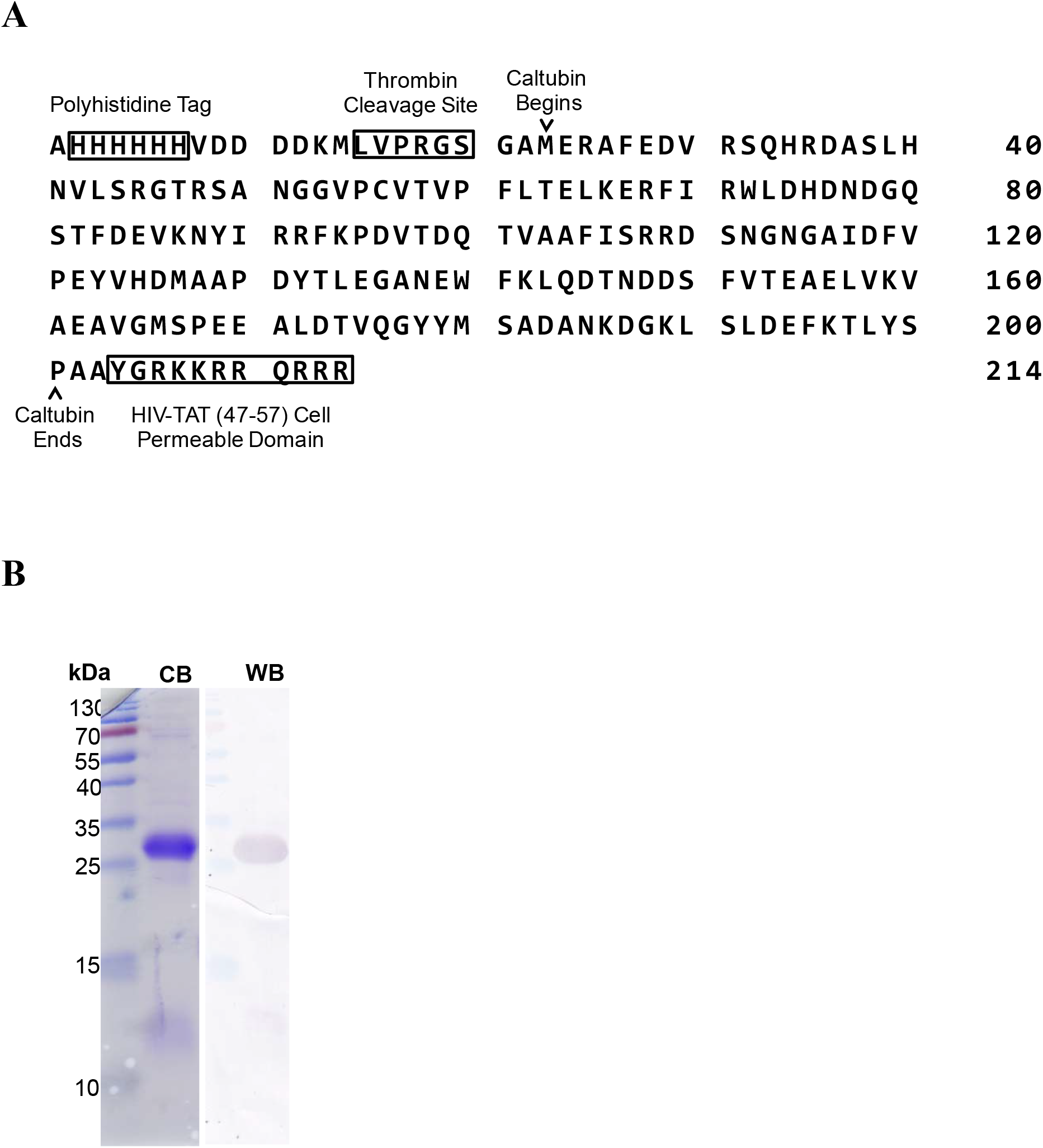
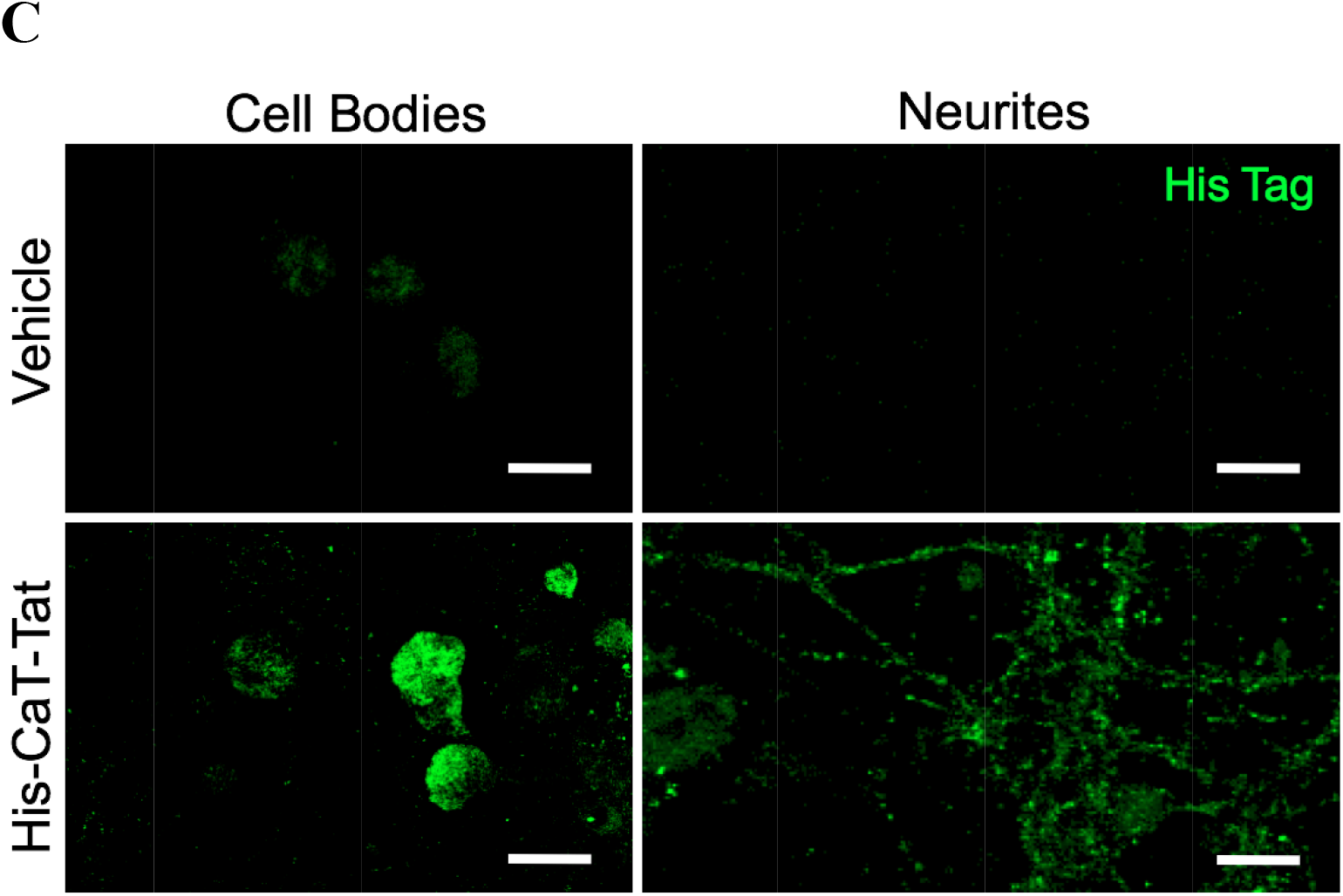
His-CaT-Tat penetrates into cultured primary hippocampal cells. (A) His-CaT- Tat protein sequence and general features. It consists of a thrombin-cleavable His6 tag to facilitate protein purification, and an arginine-rich sequence consisting of residues 47-57 of the HIV-Tat protein as the protein transduction domain. Theoretical molecular weight: 24.3 kDa. Theoretical molar absorptivity (280 nm): 20340 M^-1^cm^-1^. (B) Determination of His-CaT-Tat expression and purity: Coomassie blot (CB) showing the size and relative abundance of protein species within the sample, and a western blot (WB) depicting protein species exhibiting His6 tag immunoreactivity. (C) His-caltubin-Tat cell penetration in hippocampal culture. His-CaT-Tat cell penetration is indicated by his tag immunoreactivity in primary hippocampal culture (DIV 17) following treatment with vehicle (PBS) or 1 μM His-CaT-Tat for one hour. Cultures were washed, fixed and permeabilized prior to immunolabeling against the His6 tag and visualization with confocal fluorescence microscopy. Imaging was carried out on a narrow z-plane (small pinhole size) at a point along the z-axis corresponding to the center of the cytoplasm or neurite/neuroplasm, respectively. Scale bar represents 8 μm.

**Supplementary Figure 2.**
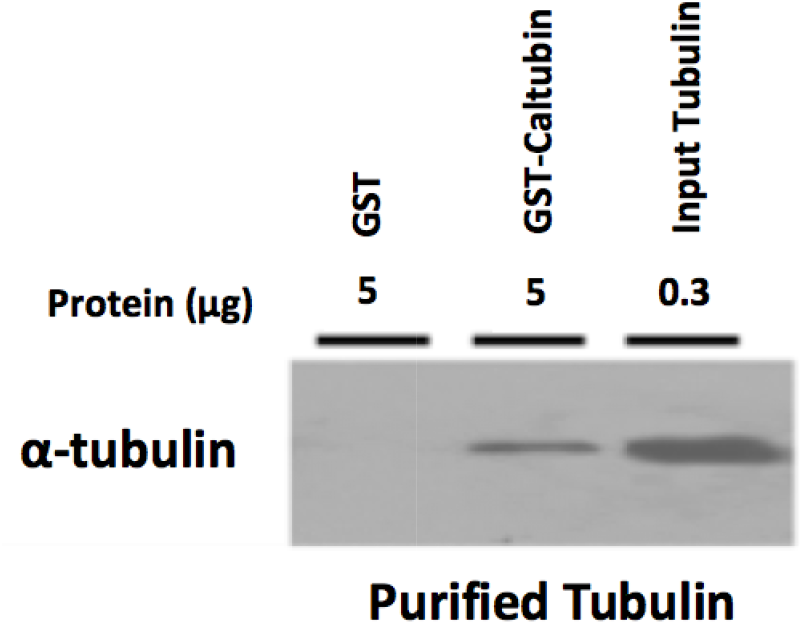
Tubulin interacts with caltubin directly. Purified α-tubulin associates with GST- caltubin more strongly than GST alone in an affinity pull-down assay. Briefly, GST or GST-caltubin was incubated with purified tubulin. The mixture was then incubated with sepharose beads to capture the GST tags and all associated proteins. Beads were then washed, and immobilized proteins were eluted prior to running a western blot and probing for α-tubulin immunoreactivity in a buffer consisting of PBS, 0.2% Triton X-100, 1 mM PMSF and protease inhibitor cocktail to determine binding.

